# An Endogenously activated antiviral state restricts SARS-CoV-2 infection in differentiated primary airway epithelial cells

**DOI:** 10.1101/2021.08.17.456707

**Authors:** Lindsay Broadbent, Connor G.G. Bamford, Guillermo Lopez Campos, Sheerien Manzoor, David Courtney, Ahlam Ali, Olivier Touzelet, Conall McCaughey, Ken Mills, Ultan F. Power

**Affiliations:** Wellcome-Wolfson Institute for Experimental Medicine, Queens University Belfast, Belfast, Northern Ireland, United Kingdom; Patrick G Johnston Centre for Cancer Research, Queen’s University Belfast, Belfast, UK; Regional Virus Laboratory, Belfast Health and Social Care Trust, Belfast, Northern Ireland

## Abstract

Severe acute respiratory syndrome coronavirus 2 (SARS-CoV-2), the cause of the coronavirus disease-19 (COVID-19) pandemic, was identified in late 2019 and went on to cause over 3.3 million deaths in 15 months. To date, targeted antiviral interventions against COVID-19 are limited. The spectrum of SARS-CoV-2 infection ranges from asymptomatic to fatal disease. However, the reasons for varying outcomes to SARS-CoV-2 infection are yet to be elucidated. Here we show that an endogenously activated interferon lambda (IFNλ) pathway leads to resistance against SARS-CoV-2 infection. Using a well-differentiated primary nasal epithelial cell (WD-PNEC) model from multiple adult donors, we discovered that susceptibility to SARS-CoV-2 infection, but not respiratory syncytial virus (RSV) infection, varied. One of four donors was resistant to SARS-CoV-2 infection. High baseline IFNλ expression levels and associated interferon stimulated genes correlated with resistance to SARS-CoV-2 infection. Inhibition of the JAK/STAT pathway in WD-PNECs with high endogenous IFNλ secretion resulted in higher SARS-CoV-2 titres. Conversely, prophylactic IFNλ treatment of WD-PNECs susceptible to infection resulted in reduced viral titres. An endogenously activated IFNλ response, possibly due to genetic differences, may be one explanation for the differences in susceptibility to SARS-CoV-2 infection in humans. Importantly, our work supports the continued exploration of IFNλ as a potential pharmaceutical against SARS-CoV-2 infection.

## Introduction

In late 2019 a novel coronavirus, severe acute respiratory syndrome coronavirus 2 (SARS-CoV-2), emerged in Hubei province in China ^1,2^. This virus is the causative agent of the coronavirus disease-19 (COVID-19) pandemic, which has resulted in over 4.2 million deaths worldwide (as of August 2021). SARS-CoV-2 can infect people of any age. However, severe disease disproportionately affects older individuals or those with underlying health conditions. Clinical manifestations of SARS-CoV-2 infection range from asymptomatic or very mild symptoms to severe illness requiring hospitalization and death. COVID-19 disease is primarily a lung disease, typically manifesting itself as acute respiratory distress syndrome in severe cases, although extra-pulmonary complications, including cardiac, vascular, kidney and neurological pathologies, are also evident ^3^. Furthermore, coagulopathies are commonly reported in COVID-19 disease patients. Reasons for the extreme variation in disease outcomes remain to be elucidated.

The upper respiratory tract is the primary site of SARS-CoV-2 infection. Infection can progress to the lower airways resulting in airway inflammation and potentially fatal pneumonia^1^. The major receptor for the viral spike protein (S), human angiotensin-converting enzyme 2 (ACE2), is expressed on airway epithelial cells but is also found on other tissue, including endothelial cells and smooth muscle cells^4^. Entry of the virus requires activation of the S glycoprotein by host proteases, such as furin and transmembrane serine protease 2 (TMPRSS2)^5^. SARS-CoV-2 is cytopathic to airway epithelial cells and alveolar cells. However, immune responses induced following infection are also thought to drive pathogenesis and disease severity in COVID-19. Chemokines and cytokines are released from infected tissues promoting leukocyte recruitment to the lungs, resulting in substantial inflammation in severe cases. Hallmarks of severe COVID-19 disease include high serum levels of IL-6, TNF-α, IP-10/CXCL10 and MCP-1, among others^6,7^.

Dependent on viral, host and environmental factors, the IFN response to infection, and subsequent inflammation, can be either beneficial or deleterious to the individual. Type I and λ (type III) interferons (IFNs) are produced in response to viral infection. IFN-mediated activation of the JAK-STAT pathway is responsible for the dramatic alteration in the cellular transcriptome and the activation of interferon-stimulated genes (ISGs), many of which have antiviral activities^8^. Patients with COVID-19 have low serum levels of type I and III IFNs but high levels of pro-inflammatory chemokines and cytokines^9^. However, it has been observed that higher serum IFN correlates positively with viral load and disease severity^10^. It is hypothesized that dysregulated, reduced or delayed IFN responses, in combination with high cytokine levels, results in more severe COVID-19 disease^11^.

Unlike type I IFN receptors, the IFNλ receptor, IFNLR1, is not ubiquitously expressed and is present predominantly on epithelial cells at barrier tissues like the lungs and gastrointestinal tract^12^. As such, type III IFNs are considered to produce a more localised response to infection in comparison to the systemic inflammatory response often induced by type I IFN^13^. SARS-CoV-2 has been shown to induce an IFN response in epithelial cells, the timing of which is delayed with regard to peak viral replication. Pre-treatment of cells with IFNs is effective at blocking SARS-CoV-2 infection, and several directly anti-SARS-CoV-2 ISGs have been discovered, such as Ly6, OAS1 and IFITMs^14–16^.

As the primary target of SARS-CoV-2 infection is the airway epithelium, models that authentically recreate the physiology and morphology of human airway epithelium *in vivo* will undoubtedly be important tools to address fundamental questions about human/SARS-CoV-2 interactions and subsequent innate immune responses. Therefore, in this study we report differential susceptibility to SARS-CoV-2 infection in well-differentiated primary nasal epithelial cell (WD-PNEC) cultures derived from adult donors. We obtained nasal brushings from 4 healthy donors with no previous history of SARS-CoV-2 infection. Interestingly, we found dramatically different susceptibility to SARS-CoV-2 infection, with 1 of the 4 donors being resistant. The resistance was likely mediated by high endogenous IFNL expression and associated ISG responses. These data may provide an explanation as to why some individuals experience asymptomatic infection following exposure to SARS-CoV-2.

## Results

### Resistance to SARS-CoV-2 infection in WD-PNECs

WD-PNEC cultures derived from 4 healthy adult donors demonstrated different susceptibility to SARS-CoV-2 infection and viral growth kinetics (Figure 1A). Following inoculation apical washes were harvested every 24 h post infection and titrated by plaque assay on Vero cells. WD-PNECs from 3 donors demonstrated similar SARS-CoV-2 growth kinetics with peak viral titres of >10^6^ PFU/mL at 48 hpi followed by a plateau or decrease at 72 hpi. Surprisingly, infection of donor 122 did not result in productive infection of SARS-CoV-2, with virus titres continually diminishing relative to titres following inoculation. Immunofluorescence reinforced these results, with high levels of SARS-CoV-2 N expression in WD-PNECs from donors 311, 43 and 32, but with very few infected cells evident in those from donor 122 (Figure 1C). Consistent with other data, SARS-CoV-2 infection was restricted to the apical surface of the cultures^17^. To determine if resistance to infection was specific to SARS-CoV-2, we infected cultures derived from the same donors with respiratory syncytial virus (RSV), a RNA virus that primarily infects ciliated airway epithelium^18^. There was no evidence of restrictions in RSV growth kinetics in any donor. (Figure 1B).

**Figure 1:**
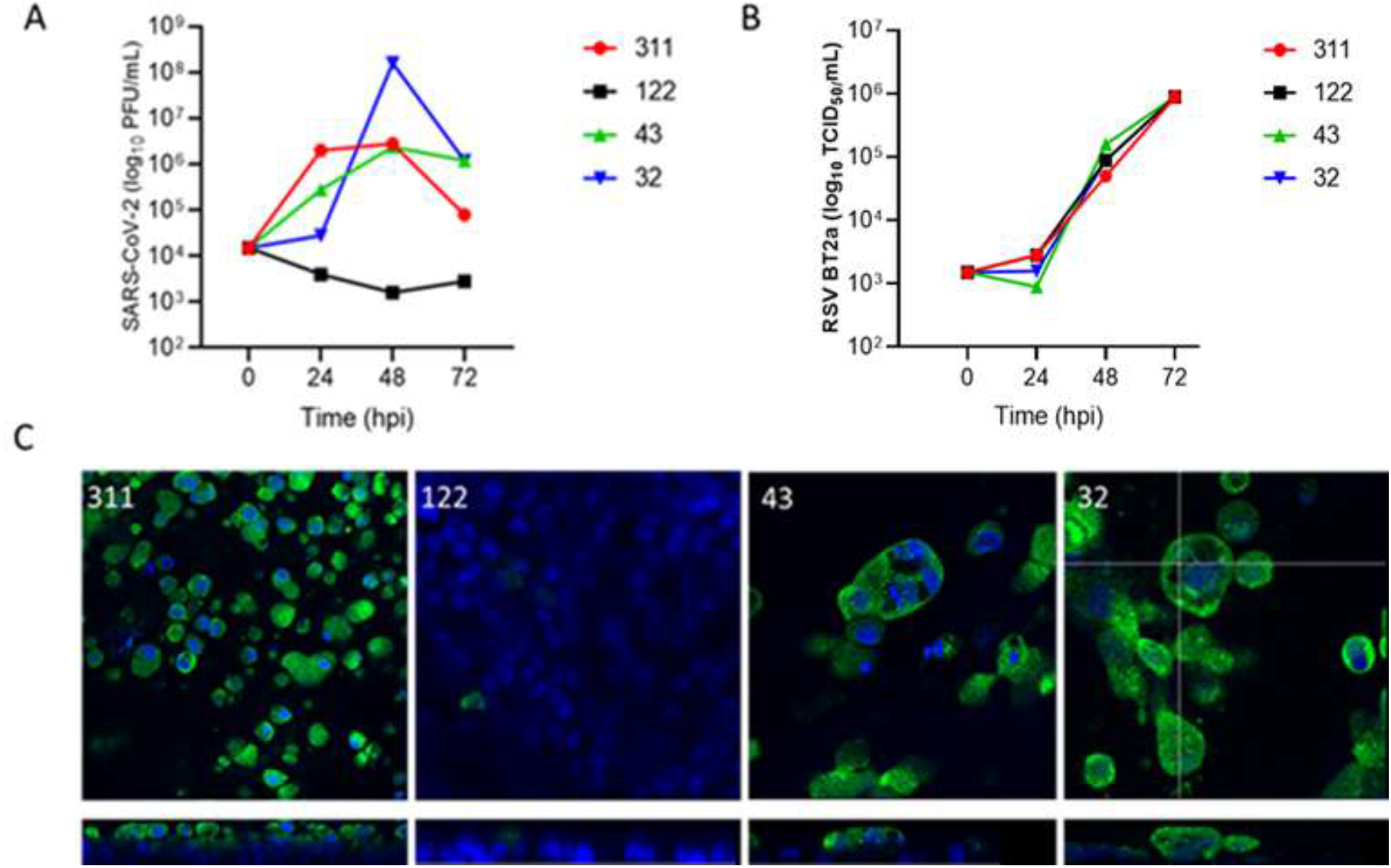
WD-PNECs from different donors are differentially susceptible to SARS-CoV-2, but not RSV. WD-PNECs from 4 adult donors were infected with a clinical isolate of SARS-CoV-2 (MOI=0.1). Apical washes were harvested every 24 h and titrated by plaque assay on Vero cells (A). Cultures were fixed at 96 hpi and SARS-CoV-2 N protein (green) was detected by immunofluorescence. Nuclei counterstained with DAPI (blue). Images and orthogonal sections were obtained on a Leica SP5 confocal microscope at x63 magnification (B). In parallel, cultures from the same donors were infected with RSV (MOI=0.1). Apical washes were harvested every 24 h and titrated on HEp-2 cells (C).

### Pre-activation of IFN pathway in resistant WD-PNECs

The timing and magnitude of IFN responses to SARS-CoV-2 infection is thought to play a role in disease severity. Transcriptomic analysis of RNA extracted at 48 and 96 hpi revealed that there was upregulation of genes associated with the IFNλ pathway in the mock-infected cultures of donor 122. Following SARS-CoV-2 infection the gene transcription signature of donor 122 did not significantly change (Figure 2A). Similar, or greater, levels of expression of *ACE2* and *TMPRSS2* were observed in this donor, confirming that resistance to infection was not due to absence or low expression of the attachment and entry factors needed for SARS-CoV-2 infection. This was confirmed by immunofluorescence (data not shown). The three donors susceptible to infection had similar patterns of gene expression, including an increase in expression of IFNλ-associated genes following SARS-CoV-2 infection (Figure 2A). To confirm these results, IFNλ1 concentrations in basolateral medium from SARS-CoV-2- or mock-infected WD-PNECs (n=4 donors) were determined at 48 and 96 hpi. Increased IFNλ1 secretion was evident following infection at 96 hpi in cultures from donors 43 and 32, but not 311. In contrast, donor 122 cultures, which were not permissive to SARS-CoV-2 infection, had high levels of IFNλ1 (> 500 pg/mL) in the basolateral medium from mock-infected cultures at 48 and 96 hpi (Figure 2B). Transcriptomic analysis of the IFIH1-mediated interferon induction and toll-like receptor (TLR) signaling pathways revealed that donor 122 had a unique gene transcription signature compared to mock or infected cultures from the other donors (Supplementary Figure 1).

**Figure 2.**
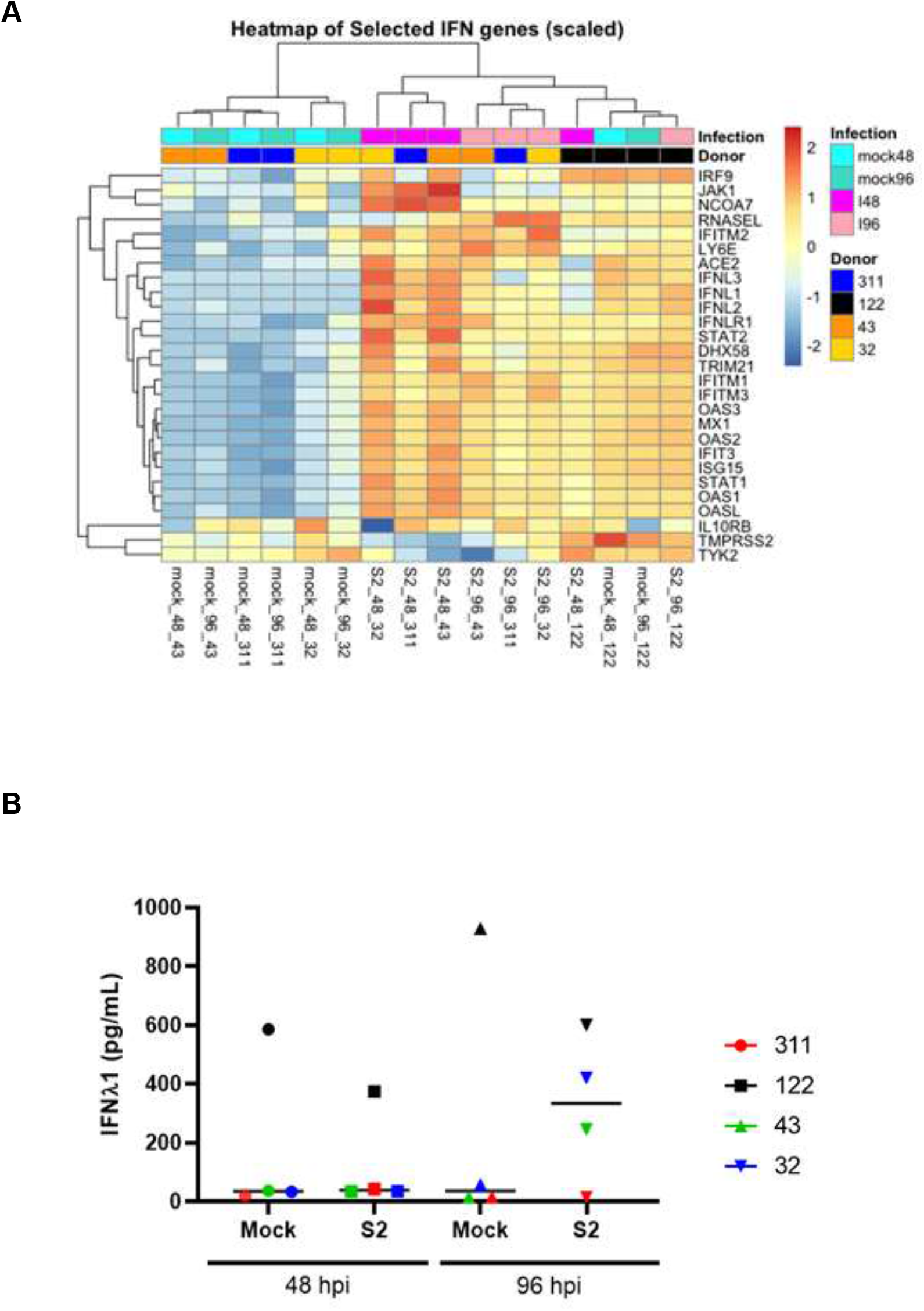
Transcriptomic profiles and protein quantification demonstrated a preactivated IFN response in a donor resistant to infection. WD-PNECs from 4 adult donors were infected with a clinical isolate of SARS-CoV-2 (MOI=0.1) or mock infected. At 48 and 96 hpi RNA was extracted from the cultures and global transcriptomic analysis was performed. Selected genes, pertaining to innate immune responses to viral infection are illustrated by heatmap, gene upregulation (Red) and down regulation (blue) (A). Preactivated immune response was validated by quantifying IFNλ1 concentration in basolateral medium from mock and SARS-CoV-2 infected WD-PNECs from 4 donors was quantified by ELISA (B).

### Pre-activation of IFN pathway and SARS-CoV-2 resistance phenotype are not transient

To determine if the high baseline level of IFNλ1 in donor 122 cultures was an artifact of culture or the time at which the nasal brush was obtained, a second brush was taken 4 months after the first and cultured in the same manner. When fully differentiated the second cultures were infected with SARS-CoV-2, with similar results (Figure 3A). WD-PNECs cultured from the second nasal brush of donor 122 also had high endogenous levels of IFNλ (∼400 pg/mL), albeit reduced compared to those evident in the cultures from the first brush. In contrast, IFNλ levels from mock-infected cultures from 6 other donors demonstrated background levels of secretion (Figure 3B).

**Figure 3.**
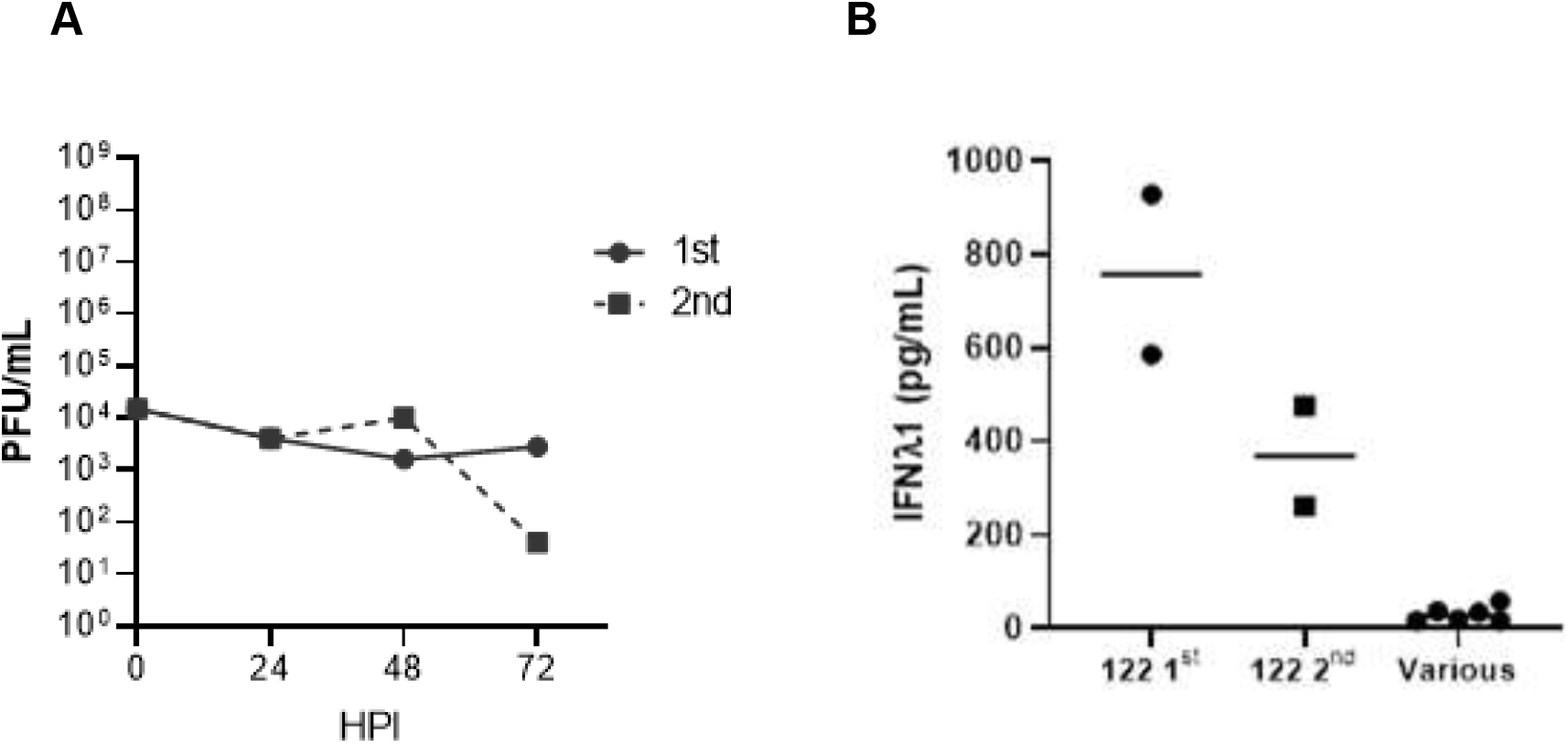
Resistance to SARS-CoV-2 infection and preactivated interferon pathway was not transient. Independent nasal brushes were obtained from donor 122 four months apart, designated 1^st^ brush and 2^nd^ brush (1^st^ brush results from experiment 1A). Cells were cultured for a minimum of 21 days in ALI until fully differentiated then infected with SARS-CoV-2 (MOI=0.1). Apical washes were harvested every 24 hpi up to 72 hpi and titrated by plaque assay. (A). IFNλ concentration in basaolateral medium from uninfected WD-PNECs from the 1^st^ and 2^nd^ nasal brushes from donor 122 were determined by ELISA. This is compared to other uninfected WD-PNEC cultures derived from 6 donors (B).

### IFN signaling restricts SARS-CoV-2 infection

SARS-CoV-2 is sensitive to exogenous IFN treatment ^19^. We hypothesised that the high baseline level of IFNλ in basolateral medium of donor 122-derived WD-PNECs limited the spread of SARS-CoV-2 within the culture and prevented productive infection. Therefore, we blocked the major signalling pathway of IFNλ with the JAK1/2 inhibitor ruxolitinib prior to infection. WD-PNECs derived from a donor with low IFNλ secretion in response to SARS-CoV-2 infection (donor 311) was used as a control. Ruxolitinib treatment resulted in a massive increase in SARS-CoV-2 titres in apical washes from cultures derived from donor 122 (>3 log10 PFU/mL), reaching a peak titre of 7.6 log10 PFU/mL at 72 hpi (Fig. 4A). As expected, there was no significant effect of ruxolitinib treatment on donor 311, which also reached peak viral titres at 72 hpi. Viral titres at 72 hpi were comparable for untreated donor 311 cultures and ruxilitinib-treated donor 122 cultures, 7.78 and 7.6 log10 PFU/mL, respectively.

**Figure 4.**
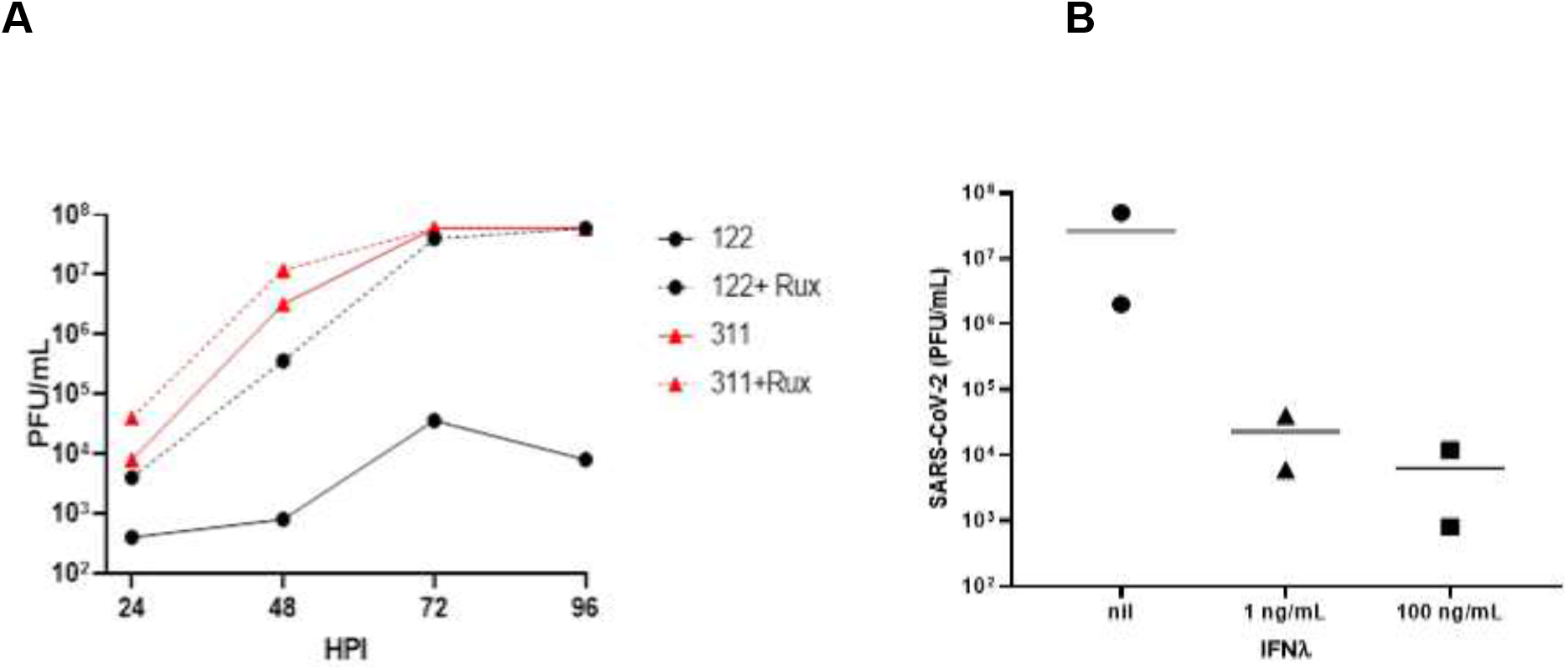
SARS-CoV-2 growth kinetics are dependent on IFN signalling. WD-PNECs derived from donor 122 and a permissive donor were treated basolaterally with 1 µM ruxolitinib for 48 h, or left untreated, prior to infection with SARS-CoV-2 (MOI=0.1). Ruxolitinib was replaced at 1 and 48 hpi. Apical washes were harvested every 24 h post infection until 96 hpi. SARS-CoV-2 viral growth kinetics were determined by plaque assay on Vero cells (A). WD-PNECs derived from a SARS-CoV-2 permissive donor were treated basolaterally in duplicate with 0, 1 ng or 100 ng/mL IFNλ for 24 h prior to infection with SARS-CoV-2 (MOI=0.1). Apical washes were harvested at 48 hpi and titrated by plaque assay on Vero cells.

To demonstrate that IFNλ was the underlying cause of resistance to SARS-CoV-2 infection we replicated these conditions in the SARS-CoV-2 permissive donor 43. WD-PNECs were pre-treated with IFNλ in the basolateral medium for 24 h prior to infection (Figure 4B). Treatment with both 1 and 100 ng/mL IFNλ greatly reduced viral titres at 48 hpi by >2 log10 PFU/mL, resulting in titres similar to the input inoculum (4.18 log10 PFU/mL).

### Single nucleotide polymorphisms (SNP) analysis revealed differences in pathogen sensing and interferon induction pathways unique to the resistant donor

Unbiased SNP analysis from our RNAseq data demonstrated clear differences between donors, with unique variants in several genes in the SARS-CoV-2-resistant donor compared to the permissive donors. Focusing our SNP analysis to selected genes associated with antiviral responses, we identified 388 SNPs unique to donor 122 and a further 33 present in donors 311, 43 and 32 but not detected in donor 122 (Figure 5A). Further conditional filtering for quality and existing SNP (according to dbSNP, NIH) identified 10 SNPs potentially involved in the high endogenous IFN secretion and resistance to SARs-CoV-2 demonstrated by WD-PNECs derived from donor 122 (Figure 5B). Interestingly, SNPs unique to donor 122 were found in ADAR (rs3738032), MAVS (rs2089960995), TLR1 (rs4833095), TLR2 (rs1816702; rs3804099) and TLR3 (rs3775296; rs3775291). SNPs present in all donors except 122 were found in ADAR (rs1127313; rs1127326) and IFIH1 (rs1990760). Together these results show that resistant versus permissive donors have unique constellations of SNPs that may cause the molecular mechanisms of susceptibility or resistance to SARS-CoV-2 infection.

**Figure 5.**
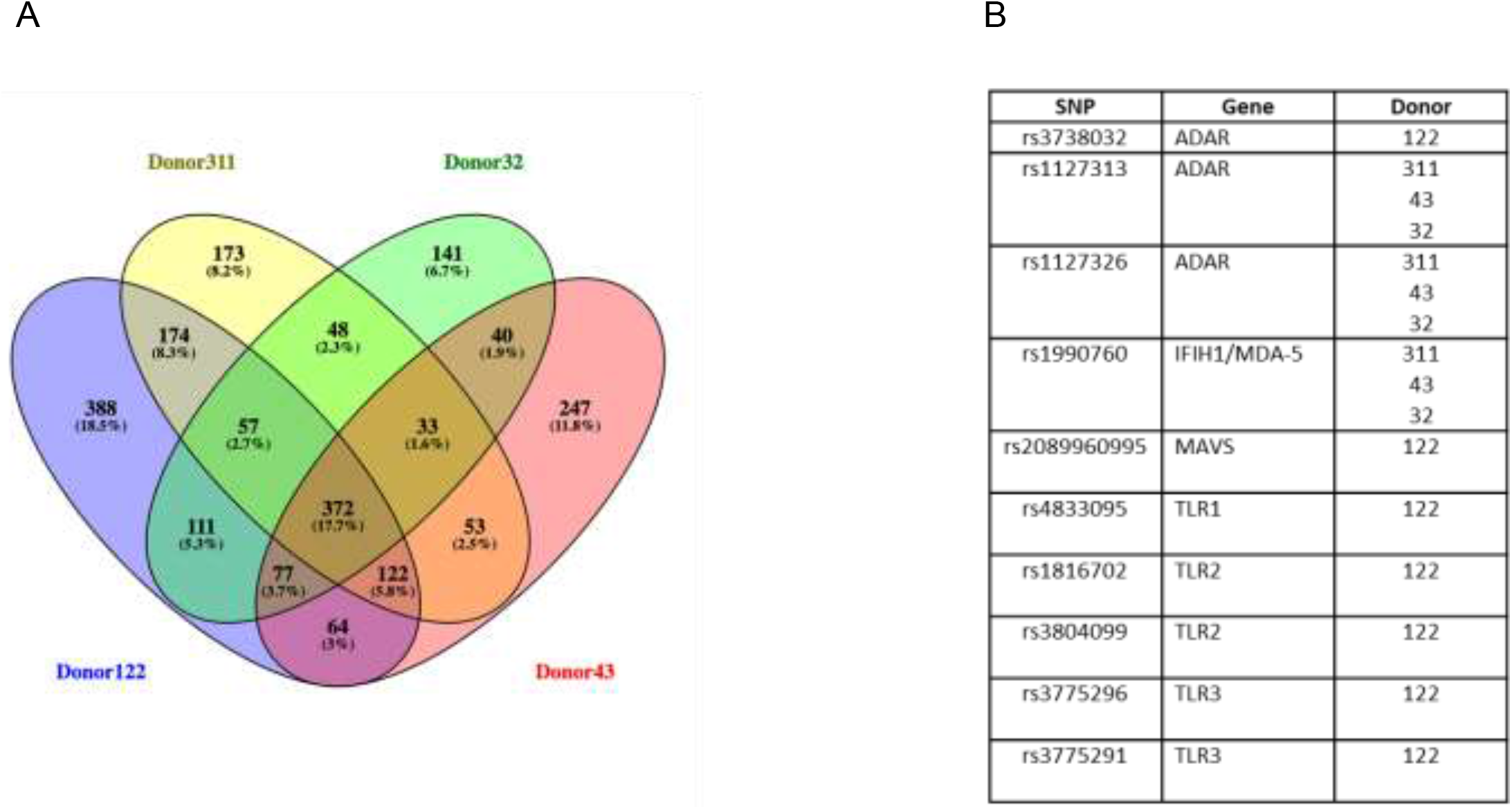
SNP analysis from transcriptomics data. SNPs unique to each donor and common displayed as a Venn diagram (A). SNPs unique to donor 122 or not present in donor 122 were filtered by quality (>10), with an existing entry in dbSNP (NIH) and previously published (B).

## Discussion

The WD-PNEC model presents an ideal opportunity to study SARS-CoV-2 infection in a clinically relevant system. The upper respiratory epithelium is thought to be the initial site of infection for SARS-CoV-2^20^. As such, a comprehensive understanding of SARS-CoV-2 infection of nasal epithelium is critical to understanding the early events of the infection process, as they are likely influential in subsequent pathogenesis. Why some patients with COVID-19 experience no or mild symptoms consistent with an upper respiratory tract infection, while others experience much more severe symptoms often resulting in acute respiratory distress syndrome (ARDS) remains to be determined. Consistent with earlier reports, we demonstrated that differentiated nasal epithelial cells can be productively infected with SARS-CoV-2 and that infection is primarily restricted to the apical surface. Surprisingly, however, we provide robust evidence of differential susceptibility of airway epithelium to SARS-CoV-2 infection, with airway epithelium from some individuals being highly susceptible and others resistant. Our data suggest that resistance is likely mediated in part by type III IFNs. Thus, we provide a potential explanation as to why some individuals may be able to control initial SARS-CoV-2 infection.

The household secondary attack rate for SARS-CoV-2 is estimated to be 16.6%, higher than that of SARS-CoV (7.5%) and MERS (4.7%)^21^. While this demonstrates that household contacts play a significant role in onward transmission of the virus it also highlights that not everyone, despite close contact with an infected individual, will be infected with SARS-CoV-2. Studies on pre-existing immunity to SARS-CoV-2 infections have focused on aspects of the adaptive immune system and have demonstrated reactive T cells in SARS-CoV-2-naive patients being important, possibly due to prior infections with an endemic coronavirus^22^. However, differences in susceptibility to infection due to innate immunity have not been reported previously. Interestingly, a recent pre-print demonstrates a link between inter-individual differences in the expression of anti-SARS-CoV-2 alleles of OAS1 in determine IFN-mediated cellular resistance to infection and subsequent disease^23^. However, in our system, we observed enhanced endogenous secretion of IFN and downstream antiviral ISGs. We propose that resistance to SARS-CoV-2 infection is, in part, due to endogenous secretion and signaling of relatively high levels type III IFN in the mucosae independent of SARS-CoV-2 infection. This fits with recent work showing high sensitivity of SARS-CoV-2 to prophylactic IFN treatment. SARS-CoV-2 sensitivity to type I and III IFN treatment was previously reported. However, contrary to our findings, these authors did not observe an interferon response to SARS-CoV-2 infection of primary human airway epithelial cells^17^. A study by Dee et al. demonstrated that rhinovirus infection induced an IFN response that blocked subsequent SARS-CoV-2 infection. They proposed that given the prevalence of rhinovirus infections within the population, it could impact the number of COVID-19 cases^24^. In our model, the underlying cause of such IFN secretion remains to be elucidated. Although this phenomenon has only been observed in nasal cells from one donor it provides an exciting precedent to explain resistance to SARS-CoV-2 infection. A large-scale study to determine the frequency of endogenously-activated IFN responses may help to further elucidate and explain differential susceptibility to COVID-19 disease in humans.

WD-PNECs that were resistant to SARS-CoV-2 infection were derived from a donor with no known history of SARS-CoV-2 infection, was unvaccinated against COVID-19 at the time of both nasal brush samplings, and was otherwise in good health. We cannot discount the possibility that the donor had a prior bacterial or viral infection that, while still allowing the epithelial cells to grow and differentiate, resulted in IFNλ production in uninfected WD-PNEC cultures. Furthermore, cultures were grown and differentiated in the presence of antibiotics. Similarly, epigenetic changes are known to occur following viral infection. Of particular note is influenza-induced epigenetic methylation of H3K79, which may play an important role in interferon signaling^25^. That the WD-PNECs cultured from the second nasal brushing 4 months after the initial sample were also resistant to SARS-CoV-2 infection, suggests that this high endogenous level of IFNλ was not a transient response, consistent with a genetic or epigenetic mechanism. Indeed, SNP analysis highlighted differences in IFN pathway genes in the donor that was resistant to SARS-CoV-2 infection compared to those who were permissive. The presence of SNPs in Toll-like receptors (TLRs) is known to affect susceptibility to disease^26^. Here we identified several SNPs unique to donor 122 in TLR1, 2 and 3. Of particular note is a common variant (minor allele frequency of ∼0.3 globally) in TLR3 (rs3775291; C>T) which may confer resistance to HIV infection^27^. The mechanism of HIV resistance is still unknown. However, heterozygous individuals have higher cytokine mRNA levels compared to C homozygotes which could restrict viral infection^28^. This phenomenon appears to be virus specific as the same mutation actually confers susceptibility to enteroviral infection^29^.

Treatment of human CoV infection with recombinant interferon is not a new concept. Type I IFNs were tested against both SARS-CoV and MERS-CoV with promising pre-clinical results but efficacy was less convincing in clinical trials^30,31^. Early clinical trials assessing the efficacy of type I IFN treatment for COVID-19, alone or in combination with other pharmaceuticals, have initially proved promising but further data are needed to draw definitive conclusions. Similarities between type I and III IFN-induced signaling and induction of antiviral and inflammatory responses are well known. However, type III IFN effects are greatest in mucosal tissues, coincident with the tissue location of its receptor on epithelial cells. Induction of type III IFNs results in prolonged expression of ISGs^32,33^. Type III IFNs have been trialed for treatment of hepatitis C virus (HCV), and hepatitis delta virus (HDV) and, more recently, SARS-CoV-2, with positive results^34–36^. Type III IFNs, with tissue-restricted receptor expression, are potentially a much more attractive option for treatment of viral infection than the more inflammatory type I IFNs with ubiquitous receptor expression. Our data supports further investigation into IFNλ, directed towards the luminal surface of the airways, to reduce the incidence of severe COVID-19. Here we have identified donor-specific resistance to SARS-CoV-2 infection in WD-PNECs concurrent with high endogenous IFNλ, likely due to genetic differences. These findings point to a previously overlooked mechanism of resistance that could be operating on a global population level.

## Methods

### Virus cultivation and titration

The SARS-CoV-2 clinical isolate BT20.1 was isolated from a patient admitted to Royal Victoria Hospital, Belfast in June 2020. Characterisation of this strain has previously been described^37^. The virus was passaged 4 times on Vero cells, including isolation. The titres of the virus stocks and experimental samples were determined by plaque assay. In brief, Vero cells were seeded at 5×10^4^ cells/well in a 24-well cell culture plate. Twenty-four hours later a 1:10 serial dilution of the virus stock or sample was prepared and inoculated onto the Vero cells. Following 1 h incubation at 37°C a semi-solid agarose overlay was added. Vero cells were further incubated for 3 days then fixed with paraformaldehyde (final concentration 4% v/v) and stained with crystal violet (5% w/v) for plaque visualisation. The origin and characterization of the clinical isolate RSV BT2a were previously described^38^. RSV titres in biological samples were determined using HEp-2 cells, as previously described^39^.

### Well-differentiated primary nasal epithelial cell (WD-PNEC) culture, infection and treatment

Nasal epithelial cells were obtained from consented healthy adults with no known history of lung disease or previous SARS-CoV-2 infection. Culture and differentiation protocols were previously described^40,41^. In brief, cells were passaged three times in Promocell Airway Epithelial Cell Growth Medium (Promocell) then seeded onto collagen-coated Transwell supports (Corning) at 3×10^4^ cells per Transwell. After 4–6 days of submersion air-liquid interface (ALI) was initiated by removing the apical medium. Cells were differentiated using Stemcell PneumaCult ALI medium (Stemcell Technologies) supplemented with hydrocortisone and heparin. Basolateral medium was replaced every 2 days. Complete differentiation took a minimum of 21 days. WD-PNEC cultures were only used when hallmarks of excellent differentiation were evident, including extensive apical coverage with beating cilia and obvious mucus production. WD-PNECs were infected with SARS-CoV-2 (MOI=0.1) by the addition of inoculum on the apical surface and incubation for 1 h at 37°C. Cultures were treated basolaterally with 0, 1 or 100 ng/mL human IFNλ3 (R&D systems, Biotechne) for 24 h. Ruxolitinib (Selleck Chemicals) (1 µM) was added to the basolateral medium 48 h before infection and was replaced at 1 and 48 hpi. Apical washes were harvested by adding 200 µL DMEM (low glucose; Gibco), incubation for 5 mins and gentle aspiration without damaging the cultures.

### Immunofluorescence

Cultures were fixed with 4% paraformaldehyde (w/v) for a minimum of 1 hour. Cells were permeabilised with 0.2% Triton X-100 (v/v) then blocked using 0.4% bovine serum albumin (BSA) (w/v). Antibodies used are specified in figure legend. Cells were counterstained with DAPI-mounting medium (Vectashield). Images were obtained on a Nikon Eclipse TE-2000U or a Leica SP5 confocal microscope.

### Sequencing and Transcriptomic analysis

WD-PNECs derived from 4 donors were infected with SARS-CoV-2 (MOI=0.1) or mock infected. RNA was extracted at 48 and 96 h for each condition, in duplicate. Cultures were incubated with 400 µl of Lysis/Binding buffer (High Pure Isolation Kit; Roche) for 5 min at RT. Total Lysed samples were vortexed for 15 sec and transferred to a High Pure filter tube. RNA was extracted as per the manufacturer’s instructions. RNA sequencing was performed via Kapa RNA HyperPrep with RiboErase (Roche). Raw sequence reads were quality checked with FastQC and pre-processed for the sequence cleaning and adapter removal using cutadapt 2.1 and Trim Galore! To remove adapter contamination. (https://www.bioinformatics.babraham.ac.uk/projects/trim_galore/). These pre-processed fastq files were then aligned to the human genome GRCh38v33 using STAR aligner version 2.7.3a^42^ using default parameters and subsequently genes were quantified using htseq version 0.11.3^43^.

Quantified genes (raw counts) were processed using edgeR^44^ for normalisation purposes and transformed into logCPMs (log counts per million). These logCPM values were subsequently used to generate a heatmap using pheatmap package for comparison and visualisation purposes using a selection of interferon related genes extracted from a selection of pathways annotated in WikiPathways and Reactome Pathway Database (WP2113; R-HSA-913531; WP1904; R-HSA-168928; WP4880; WP1835; R-HSA-909733; WP75), ACE2 and TMPRSS2. SNP calling was carried out using ExactSNP program from the subread package using default parameters^45^. To increase the sequencing depth, we merged together all samples from each donor and used the merged file as input for the SNP calling phase. In addition to SNP calling we also annotated the variants identified using SnpEff software and the GRCh38.99 database^46^.

### ELISA analysis

Concentrations of IFN-λ1 in basolateral medium samples from WD-PNECs were measured using a human IFN-λ1/IL-29 ELISA (Invitrogen) according to the manufacturer’s instructions.

## Acknowledgements

The authors would like to thank the following individuals and groups for their contribution and assistance: QUB FMHLS Genomics Core Technology Unit; QUB staff Grace Roberts, Judit Barabas, Mervyn McCaigue, Cathy Fenning, Nuala McCann and David Norwood for their guidance and assistance with biological safety and maintenance of the BSL3 laboratory.

