## Supplemental Figure 1 for "An Endogenously activated antiviral state restricts SARS-CoV-2 infection in differentiated primary airway epithelial cells"

### Slide 1
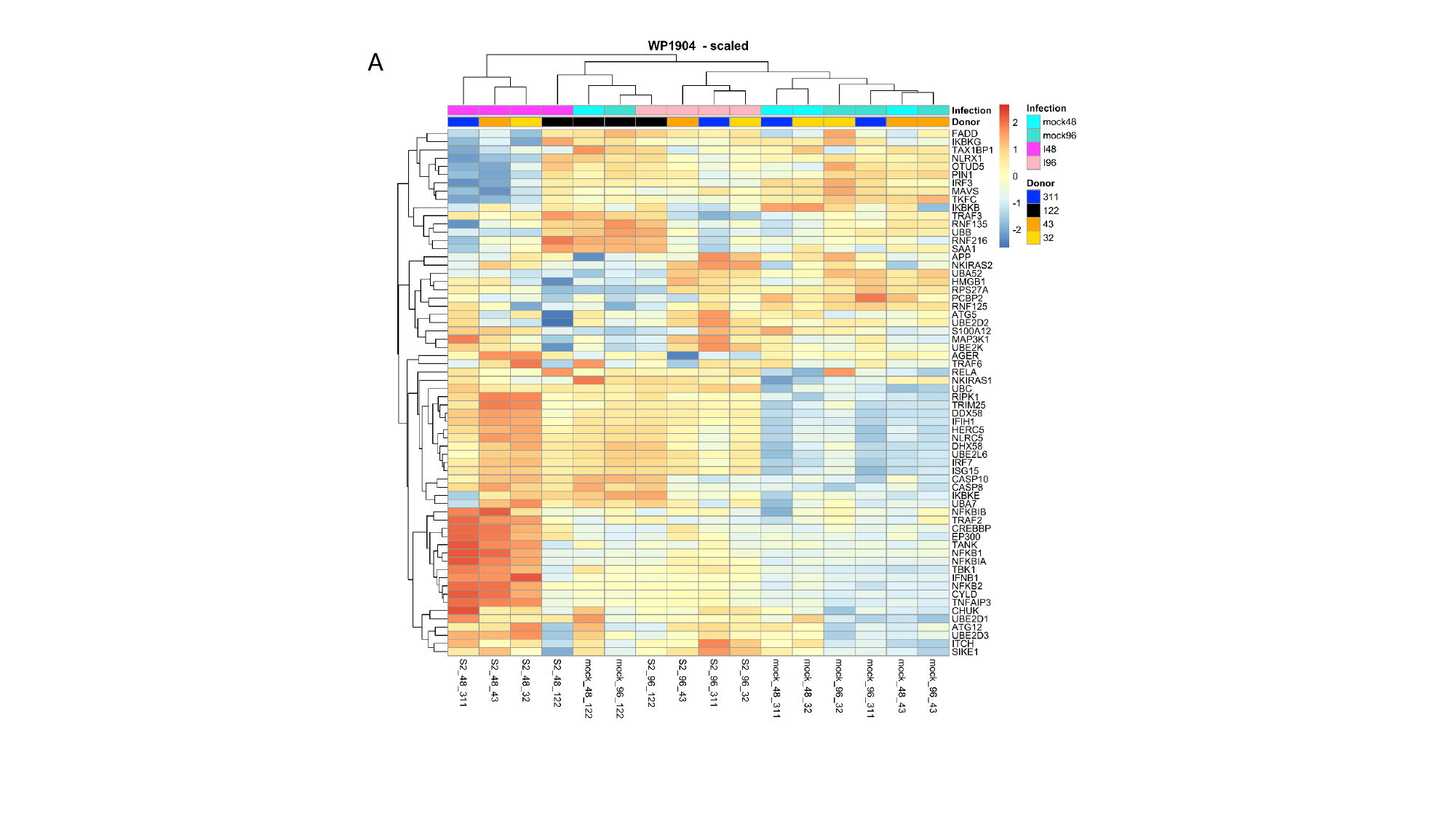

A

### Slide 2
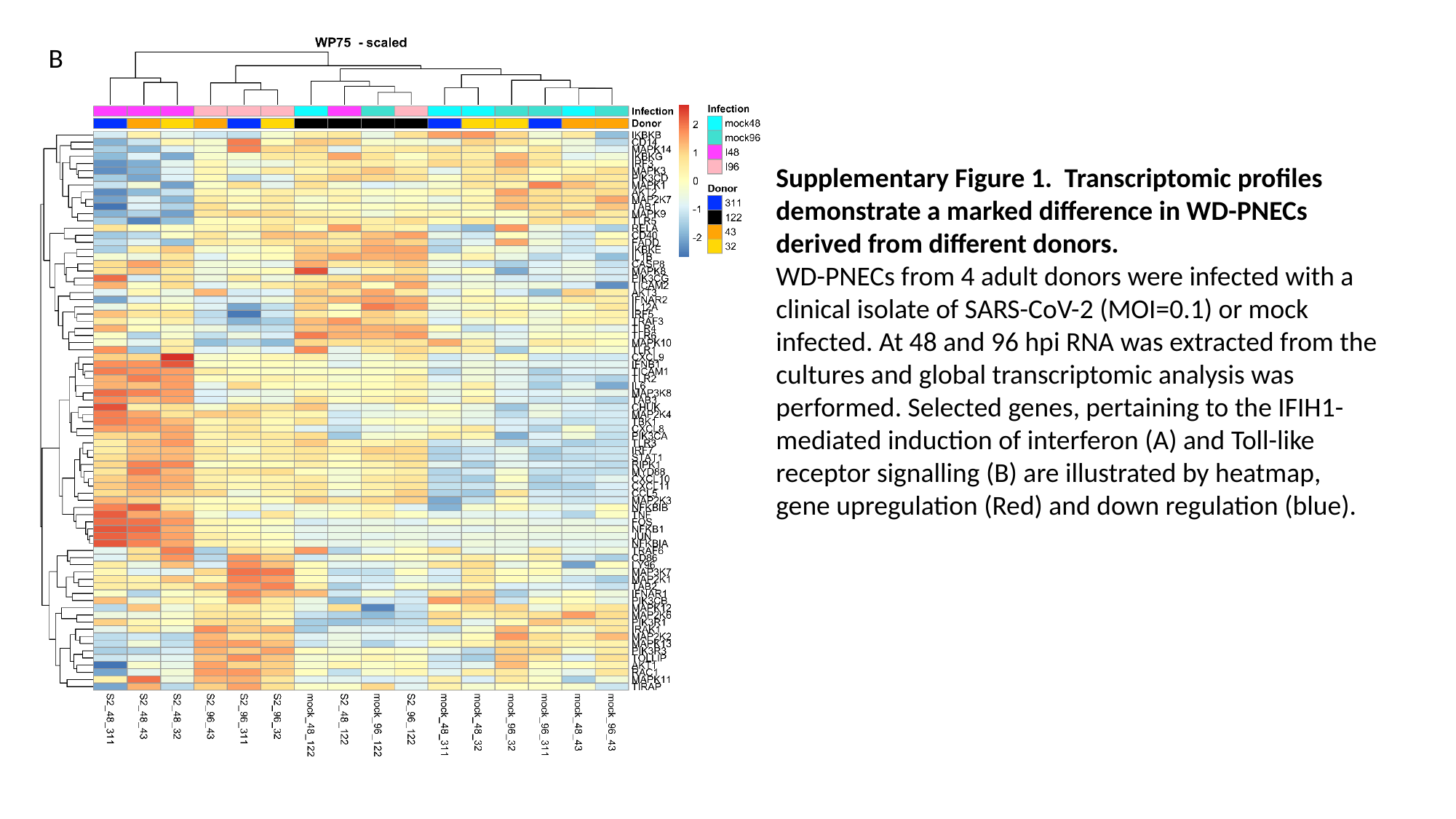

B
Supplementary Figure 1. Transcriptomic profiles demonstrate a marked difference in WD-PNECs derived from different donors.
WD-PNECs from 4 adult donors were infected with a clinical isolate of SARS-CoV-2 (MOI=0.1) or mock infected. At 48 and 96 hpi RNA was extracted from the cultures and global transcriptomic analysis was performed. Selected genes, pertaining to the IFIH1-mediated induction of interferon (A) and Toll-like receptor signalling (B) are illustrated by heatmap, gene upregulation (Red) and down regulation (blue).
